# Recurrent Duplication and Diversification of Acrosomal Fertilization Proteins in Abalone

**DOI:** 10.1101/2021.10.14.464412

**Authors:** J. A. Carlisle, M. A. Glenski, W. J. Swanson

## Abstract

Reproductive proteins mediating fertilization commonly exhibit rapid sequence diversification driven by positive selection. This pattern has been observed among nearly all taxonomic groups, including mammals, invertebrates, and plants, and is remarkable given the essential nature of the molecular interactions mediating fertilization. Gene duplication is another iimportant mechanism that facilitates the generation of molecular novelty. Following duplication, paralogs may parse ancestral gene function (subfunctionalization) or acquire new roles (neofunctionalization). However, the contributions of duplication followed by sequence diversification to the molecular diversity of gamete recognition genes has been understudied in many models of fertilization. The marine gastropod mollusk abalone is a classic model for fertilization. Its two acrosomal proteins (lysin and sp18) are ancient gene duplicates with unique gamete recognition functions. Through detailed genomic and bioinformatic analyses we show how duplication events followed by sequence diversification has played an ongoing role in the evolution of abalone acrosomal proteins. The common ancestor of abalone had four members of its acrosomal protein family in a tandem gene array that repeatedly experienced positive selection. We find that both sp18 paralogs contain positively selected sites located in different regions of the paralogs, consistent with a subfunctionalization model where selection acted upon distinct binding interfaces in each paralog. Further, a more recent species-specific duplication of both lysin and sp18 in the European abalone *H. tuberculata* is described. Despite clade-specific acrosomal protein paralogs, there are no concomitant duplications of egg coat proteins in *H. tuberculata*, indicating that duplication of egg proteins *per se* is not responsible for retention of duplicated acrosomal proteins. We hypothesize that, in a manner analogous to host/pathogen evolution, sperm proteins are selected for increased diversity through extensive sequence divergence and recurrent duplication driven by conflict mechanisms.

## Introduction

Despite their essential role in many organisms, genes functioning in fertilization or sexual reproduction are often rapidly diverging between closely related species including mammals, birds, fish, and invertebrates (Carlisle & Swanson, 2020; Swanson, Nielsen, & Yang, 2003; Swanson & Vacquier, 2002). Some pairs of interacting sperm and egg gamete recognition proteins have been shown to be rapidly co-evolving, pointing to sexual conflict or sexual selection driving the rapid evolution of fertilization genes (Bianchi, Doe, Goulding, & Wright, 2014; Clark et al., 2009; Grayson, 2015; Kamei & Glabe, 2003). This rapid diversification of sperm and egg gamete recognition proteins at a sequence level can result in species-specific fertilization function (Avella, Baibakov, & Dean, 2014; Raj et al., 2017; Zigler, McCartney, Levitan, & Lessios, 2005). Investigations into the evolution of reproductive genes paired with characterization of species-specific function can provide unique insights into infertility and reproductive isolation (Lehmann, 2018). In addition to sequence evolution, gene duplication can also contribute to the molecular diversification of reproductive protein families. *Drosophila* seminal fluid proteins often undergo duplication and diversification, with many of these duplications being species-specific (Almeida & Desalle, 2009; Doty, Wilburn, Bowen, Feldhoff, & Feldhoff, 2016; Findlay, Yi, Maccoss, & Swanson, 2008; Sirot et al., 2014; Wagstaff & Begun, 2005; D. B. Wilburn, Arnold, J. A., Houck, L. D., Feldhoff, P. W., Feldhoff, R. C., 2017). Mammalian genes mediating fertilization including CatSper, Izumo, and the ZP gene families also demonstrate interesting patterns of duplication (Aagaard, Vacquier, MacCoss, & Swanson, 2010; Cai & Clapham, 2008; Cooper & Phadnis, 2017; Grayson & Civetta, 2012). Here, we investigate how both sequence diversification and duplication together contribute to the molecular diversification of fertilization proteins.

The marine gastropod abalone (Genus *Haliotis*) is a classic model system for studying the function of gamete recognition proteins and their evolution. Abalone sperm have an extremely large acrosome containing two gamete recognition proteins, lysin and sp18 (Lewis, Leighton, & Vacquier, 1980). Lysin is the sperm mediator of the dissolution of the egg’s vitelline envelope (VE), and sp18 is thought to mediate sperm-egg plasma membrane fusion (Carlisle & Swanson, 2020; D. B. Wilburn, Tuttle, Klevit, & Swanson, 2018). The abalone egg VE is an elevated glycoproteinaceous layer homologous to the mammalian egg zona pellucida (ZP) (Carlisle & Swanson, 2020). The abalone VE and mammalian ZP are biochemically and structurally similar (Mozingo, Vacquier, & Chandler, 1995) and both contain proteins with ZP-N domains (Avella et al., 2014; Carlisle & Swanson, 2020). Binding between lysin and the ZP-N domains of the vitelline envelope receptor for lysin (VERL) leads to the non-enzymatic dissolution of the VE (Aagaard, Springer, Soelberg, & Swanson, 2013; Raj et al., 2017; Swanson & Vacquier, 1997). Three-dimensional structures of lysin and VERL have been investigated using NMR and crystallography and their ability to bind each other have been quantified (Aagaard et al., 2013; Kresge, Vacquier, & Stout, 2000, 2001; Raj et al., 2017; D. B. Wilburn et al., 2018). The highly fusogenic protein sp18 is the putative mediator of sperm-egg plasma membrane fusion in abalone (Kresge et al., 2001; Swanson & Vacquier, 1995). The structure of sp18 has been determined via crystallography, but its binding receptor of is unknown (Kresge et al., 2001).

Investigations into the evolution of abalone reproductive proteins have provided valuable insights within the field of reproductive biology. Lysin, sp18 and VERL have each been shown to evolve under positive selection by analysis of the ratio of rates of nonsynonymous substitutions to rates of synonymous substitutions (*dN/dS* > 1) (Galindo, Vacquier, & Swanson, 2003; Lee, Ota, & Vacquier, 1995; Swanson & Vacquier, 1995). Further, population genetic analysis indicate that lysin and its binding partner VERL are coevolving with each other (Clark et al., 2009). Previous studies of abalone egg and sperm gamete recognition proteins hint at the importance of duplication events for their evolution. Despite their divergent functions in reproduction and low sequence similarity, sp18 and lysin are paralogs with similar three-dimensional protein structures (Kresge et al., 2001). The common ancestor of lysin and sp18 is hypothesized to have mediated both VE dissolution and sperm egg fusion, then parsed these combined ancestral functions post-duplication to lysin and sp18, respectively.

Additional evidence suggests that duplication may be contributing to the evolution of abalone sperm fertilization proteins on a more recent timescale. The European abalone *H. tuberculata* has a species-specific duplication of lysin (Clark, Findlay, Yi, MacCoss, & Swanson, 2007). On the egg side there is also evidence of extensive gene duplication. In addition to VERL, abalone VEs contain ∼30 homologous VEZP proteins containing ZP-N domains (Aagaard et al., 2010; Aagaard, Yi, MacCoss, & Swanson, 2006). Many of these VEZPs may be structural components of the VE that play no role in gamete recognition (Killingbeck & Swanson, 2018). However, one protein (VEZP-14) is a close paralog of VERL that has undergone positive selection and is capable of binding lysin (Aagaard et al., 2013). Abalone VEZPs have only been described in two species (*H. rufescens*, and *H. fulgens*) from the North American clade. It is unknown whether there is variation in gene content across more distantly related abalone species (Aagaard et al., 2006).

In this study we investigated the contributions of duplication and sequence diversification to the evolution of proteins mediating fertilization across the genus *Haliotis*. Using new testes and ovary transcriptomic data and published genome assemblies we discovered novel duplications of the acrosomal proteins lysin and sp18. Some of these paralogs are ancestral to abalone and others are clade-specific. Further we discover signatures of positive selection in many of the paralogs and identify patterns of positively selected sites indicative of subfunctionalization. Our detailed evolutionary genomic analysis reveals how recurrent patterns of duplication paired with diversification led to the evolution of abalone gamete recognition proteins and their variation between species. Repeated duplications within the protein family containing lysin and sp18 parallels the duplication and diversification of other reproductive protein families, such as mammalian Izumo and Catsper families.

## Materials and Methods

### PacBio Library Preparation and Sequencing

To identify potential transcripts present in abalone gonadal tissue, methods were adapted from the PacBio Iso-seq protocolto create cDNA libraries. Ovary and testes transcriptome libraries were prepared for PacBio sequencing. RNA was extracted from *H. tuberculat*a ovary and testes samples by cesium chloride density gradient centrifugation (MacDonald, Swift, Przybyla, & Chirgwin, 1987). RNA samples were enriched for mRNA by using the Oligotext mRNA Mini Kit from Qiagen. Purified mRNA was used as the template for single stranded cDNA synthesis using the Clontech SMARTer cDNA Synthesis Kit. The cDNA was amplified by PCR using the AccuPrime High-Fidelity *Taq* system (Invitrogen, Carlsbad, CA, USA). Double stranded synthesis conditions were optimized with the following PCR program: °C for 2 minutes, followed by 20 cycles of 94°C for 30 s, 55 °C for 30 s, and 68 °C for 10 minutes. We used unique identifying barcoded PCR primers to amplify testes (barcode: CTGCGTGCTCTACGAC) and ovary (barcode: TCAGACGATGCGTCAT) cDNA. Because of size bias during PacBio sequencing, double stranded cDNA was fractionated using Ampure XP Beads (Beckman Coulter Life Sciences) into two fractions (Ratio of 4:1 of 0.45x : 0.6x size selection). Testis and ovary cDNA were sequenced using the PacBio RSII and was performed by the Washington State University Genomics Core.

### Identification of Acrosomal Protein Paralogs

Sequences of sp18 and lysin from the genus *Haliotis* were retrieved from NCBI Genbank sequence repository (accession numbers for lysin: L26270-79, L26281-83, L35180-81, L36589, M34388-89, M59968-72, M98874-75, HM582239; accession numbers for sp18: L36552-54, L36589-90, MN102340-42). These sp18 and lysin sequences were used as the initial query sequences when identifying paralogs in abalone transcriptomes and genomes. Queries of the *H. rufescens* Illumina-based testis transcriptome (Palmer et al., 2013) and the *H. tuberculata* PacBio testes transcriptome were conducted with tblastn with an e-value cutoff of 1e-10. Significant matches from the testes transcriptomes were searched against the NCBI sequence repository (July 2020) using tblastn in order to confirm homology to lysin or sp18 (McGinnis & Madden, 2004). New sequences were uploaded to Genbank under accession numbers OK491874-OK491877.

Regions of publicly available abalone genomes containing novel acrosomal protein duplications of sp18 and lysin were identified by using tblastn with a e-value cutoff of 1e-10 (Botwright et al., 2019; Gan et al., 2019; Masonbrink et al., 2019; Nam et al., 2017). Samtools faidx was used to extract the region of scaffolds containing the tblastn hits and 20,000 base pairs upstream and downstream of the hit. We predicted the exonic sequences of the sp18 and lysin paralogs from these extracted regions using the Protein2Genome command of the program Exonerate version 2.2.0 (Slater & Birney, 2005). The top scoring prediction from Exonerate was used to define the paralog’s exons. We used the same lysin and sp18 sequences from the tblastn search as query sequences. For all full-length sequences, the SignalP-5.0 prediction server was used to predict presence of functional signal peptides (Almagro Armenteros et al., 2019). Presence of signal peptides were predicted with probabilities >0.9; the signal peptide cleavage site was predicted with a probability >0.5.

### Phylogenetic Analysis

The phylogenetic inference tool RAxML-NG was used to construct all phylogenetic trees with the LG substitution matrix (Kozlov, Darriba, Flouri, Morel, & Stamatakis, 2019; Le & Gascuel, 2008). RaxML-NG conducts maximum likelihood based phylogenetic inference and provides branch support using non-parametric bootstrapping (Kozlov et al., 2019). The best scoring topology of 20 starting trees (10 random and 10 parsimony-based) was chosen. RaxML-NG was used to perform non-parametric bootstrapping with 1000 re-samplings that were used to re-infer a tree for each bootstrap replicate MSA. Finally, we mapped the bootstrap scores on the best-scoring starting tree. The Transfer Bootstrap Expectation (TBE) was used as a branch support metric (Lemoine et al., 2018).

DNA multiple sequence alignments (MSA) for phylogenies of lysin and sp18 and their respective paralogs were constructed. First the protein sequences of the genes were aligned using PROMALS3D (Pei, Kim, & Grishin, 2008; Pei, Tang, & Grishin, 2008). PROMALS3D uses protein three-dimensional protein structures to inform protein alignments, and representative PDBS for lysin (5UTG) and sp18 (1GAK) were specified (Kresge et al., 2000; D. B. Wilburn et al., 2018). The protein MSAs were used to create DNA alignments of the same genes using the Pal2Nal sever (Suyama, Torrents, & Bork, 2006). For phylogenetic analysis of *H. rufescens* and *H. tuberculata* VEZP sequences, the protein sequences of the C-terminal ZP modules of each of the proteins were aligned using PROMALS3D (Pei, Tang, et al., 2008).

### Syntenic Comparison between *Haliotis rufescens* and *H. rubra*

The published genome of Haliotis rufescens is annotated with ORFs identified via transcriptomic sequencing (Masonbrink et al., 2019). We collected the sequences of 2-3 large annotated ORFs surrounding lysin, sp18, and their newly described paralogs within the *H. rufescens* genome. We used these sequences as BLAST queries against the *H. rubra* genome (Gan et al., 2019). The top hits for the *H. rufescens* ORFs were annotated onto the *H. rubra* genome and used to establish synteny *between H. rubr*a scaffold 62 and the *H. rufescens* scaffolds 48 and 101. A reciprocal blast of the regions identified as orthologous ORFs in *H. rubra* were queried against the *H. rufescens* genome to verify orthology.

### Detecting Selection and Positively Selected Sites

Values of *d*_*N*_*/d*_*S*_ for genes were estimated using the codeml program of PAML 4.8 (Yang, 2007). We compared models of selection using a likelihood ratio test (LRT) between neutral models and models with positive selection. Specifically, we compared M1 v. M2, M7 v. M8, and M8a v. M8. Likelihood ratio tests were performed where the likelihood ratio (LRT) statistic was twice the negative difference in likelihoods between nested models. For M1a v M2a or M7 v Model 8 the LRT was compared to the χ^2^ distribution with 2 degrees of freedom (Yang, 2007). For the M8a v. M8 comparison, twice the negative difference in likelihoods between the nested models being compared, the LRT statistic, is approximated by the 50-50 mixture distribution of 0 and χ^2^ with degree of freedom 1 (Swanson et al., 2003). To identify specific sites in proteins evolving under positive selection, we used Naive Empirical Bayes (Pr(omega > 1) = 0.95). The sites under positive selection were identified using Model 8.

### Testing for divergence in regions undergoing positive selection in duplicate sperm proteins

We designed three unbiased tests to determine if sites under positive selection in either sp18 or sp18-dup are differentially clustered between paralogs. First, we created a parametric test based on the Wald-Wolfowitz runs test (Magel, 1997) to determine whether positively selected sites in the paralogs sp18 and sp18-dup were non-randomly distributed in a protein alignment of both paralogs. The Wald-Wolfowitz runs test determines the randomness of a two-category data string by examining changes between categories by counting “runs.” We designed a parametric version of the test to allow the inclusion of three categories. The categories were sites under positive selection in sp18, sites under selection in sp18-dup, and sites under selection in both. The order in which these sites under selection in the categories appeared in an alignment of *H. fulgens* sp18 and *H. sorenseni* sp18-dup became our data string. For the data string generated from our paralog alignment we counted how many times the identity of sites in the string changed plus one. A visualization of the pipeline for preparing this data string and counting “runs” is shown in Supplementary Figure 1A.

To make a parametric version of this runs test, we generated 1000 simulated data strings via bootstrapping based on the proportion of sites shown to be under positive selection in either paralog. For each of these simulated data strings we also calculated the number of runs. The number of data strings with a count of “runs” less than or equal to the count of “runs” found in the data string derived from the paralog protein alignment divided by the total number of simulated data strings gives the parametric probability that by random chance categories of sites would be more or equally clustered compared to the true clustering observed. We also performed a version of the test where sites under selection in both paralogs were eliminated from the analysis. When these sites were eliminated the parametric runs test retained statistical significance (p-value < 0.001).

For our second test we evaluated whether sites under positive selection in sp18 vs sp18-dup were distributed throughout the sp18 crystal structure (1GAK) in a significantly different way. In MATLAB Online version 9.9, we identified the plane of best fit between the c-alpha carbons (the first carbon attached to the functional group of an amino acid) of the sp18 crystal structure using linear regression (Supplementary Figure 2A). This plane divided the sp18 crystal structure into two sides that we arbitrarily designated “left” and “right” (Supplementary Figure 2B). We mapped the 43 sites in sp18 and the 33 sites in sp18-dup that are under positive selection onto the sp18 crystal structure. Given the number of amino acids sites in each side of the crystal structure, we estimated the expected number of positively selected sites from each paralog that would be expected to be located on either side as the number of amino acid sites on a side divided by the total number of amino acid sites in the molecule and multiplied by the number of positively selected sites in a sp18 paralog. We used a chi-square test to examine whether the real distribution of sites between categories rejected the null expectation. This test determined whether the distribution of sites under selection in either paralog was not distributed similarly between the sides of the protein. We also created a plane perpendicular to the plane of best fit to divide the sp18 crystal structure into the categories “top” and “bottom” (Supplementary Figure 2C). We repeated an analysis for this new pair of categories that is identical to what was described previously. This analysis gave us a sense if sites under selection in either paralog were clustered nonrandomly throughout the crystal structure in different ways.

For our last clustering test we determined whether sites under positive selection in a paralog were more likely to be close in proximity in three-dimensional space to another site under positive selection in the same paralog rather than a site under positive selection in the other paralog. For each site that was under selection in a paralog, we identified the closest positively selected site in three-dimensional space that belonged to either paralog. We then calculated the expected number of times by chance the closest adjacent site for each site under positive selection would belong to the same gene rather than the other paralog. We used a chi-square test to determine whether observed sites under selection in one paralog were statistically more likely to be close to positively selected sites belonging to the same paralog than what would be expected by chance. This test has four categories of sites (for each paralog the nearest site could belong to the same paralog or not), and therefore three degrees of freedom were used to determine the p-value. Sites that were under selection in both paralogs were counted twice in this analysis since these sites were undergoing positive selection independently in both paralogs. When the closest adjacent site to a positively selected site in one paralog was undergoing positive selection in both paralogs, the adjacent site was treated as belonging to the same paralog.

### Identification of Sp18 peptides

*H. tuberculata* testis tissue was homogenized in 1% sodium dodecyl sulfate with BME at 70 °C for 30 minutes. Testis samples were separated by SDS-PAGE using a Tris-Tricine buffering system with discontinuous 4% resolving/15% separating acrylamide gels. Samples were electrophoresed at 50V for 15 min followed by 100 V for 90 min. The gel was run with the BioRad Broad Range Ladder and stained with Coomassie Blue R-250 for 15 minutes. Using the ladder as reference, the lysin and sp18-containing region (∼14-22 kDa) of the polyacrylamide gel was excised using a clean scalpel, with multiple rounds of perfusion with an ammonium bicarbonate solution followed by acetonitrile to extract detergents and salts. Trypsin proteolysis of immobilized proteins was by perfusion of a Trypsin solution (40 ug/ml stock Trypsin 1:10 in 50 mM ammonium bicarbonate) and incubation at 37°C overnight. The supernatant from the digest was collected along with the supernatant from two rounds of hydration with ammonium bicarbonate and extraction with 50% acetonitrile. The collected supernatant containing the liberated peptides was concentrated to a dry pellet using a vacuum centrifuge then reconstituted in 0.1% FA for liquid chromatography tandem mass spectrometry (LC/MS-MS). Unique peptides for sp18 copy #1 were identified in the sample using the Crux toolkit comet command (Park, Klammer, Kall, MacCoss, & Noble, 2008). The protein sequence database was composed of a six-frame translation of the *H. tuberculata* testis transcriptome.

### Identification of ZP Proteins

An exhaustive BLAST search of the *H. tuberculata* ovary transcriptome identified all cDNA sequences with homology to *H. rufescens* VEZPs. A previous study used a similar approach to originally identify known VEZPs in *H. rufescens* indicating that this approach should be sufficient to identify novel VEZPs (Aagaard et al., 2010). All cDNA sequences that matched VEZPs were filtered for duplicates using CD-HIT-EST with a threshold of 0.9 sequence identity (Huang, Niu, Gao, Fu, & Li, 2010). The longest sequence from each cluster created by CD-HIT-EST was chosen as the cluster’s representative sequence. All *H. tuberculata* sequences from this filtering process were translated and the C-terminal ZP modules were identified by identifying conserved cysteine residues. The ZP module protein sequences from both *H. tuberculata* and *H. rufescens* were aligned using PROMALS3D (Pei, Tang, et al., 2008). The MSA of these ZP modules from were used to construct a VEZP homolog protein phylogeny using the same RAXML-NG protocol described above for lysin and sp18 paralog phylogenies. New sequences were uploaded to Genbank under accession numbers OK491878-OK491909.

## Results

### Genomic Analysis Reveals Tandem Duplications of Ancestral Abalone Acrosomal Proteins

By pairing phylogenetic and genomic analysis of abalone species belonging to the North American clade (*H. rufescens, H. sorenseni, H. discus*) and the Australian clade (*H. rubra, H. laevigata*), we identifed ancestral duplications of both sp18 and lysin (**Figure 1;** sp18-dup and lysin-dup, respectively). We calculated maximum likelihood DNA phylogenies independently for lysin and sp18 with their paralogs and rooted the phylogenies by orthology (**Figure 1A & 1B**). Predicted intron/exon boundaries of the novel acrosomal protein paralogs were shared with lysin and sp18 (Metz, Robles-Sikisaka, & Vacquier, 1998). No mutations causing pseudogenization were detected within the predicted CDS of either paralog. For the abalone species with published genomes, only one (*H. rufescens*) has a published testes transcriptome (Palmer et al., 2013). Full-length sequences of lysin, sp18, and sp18-dup are expressed in the testes transcriptome of *H. rufescens*; however, lysin-dup was not detected.

**Figure 1:**
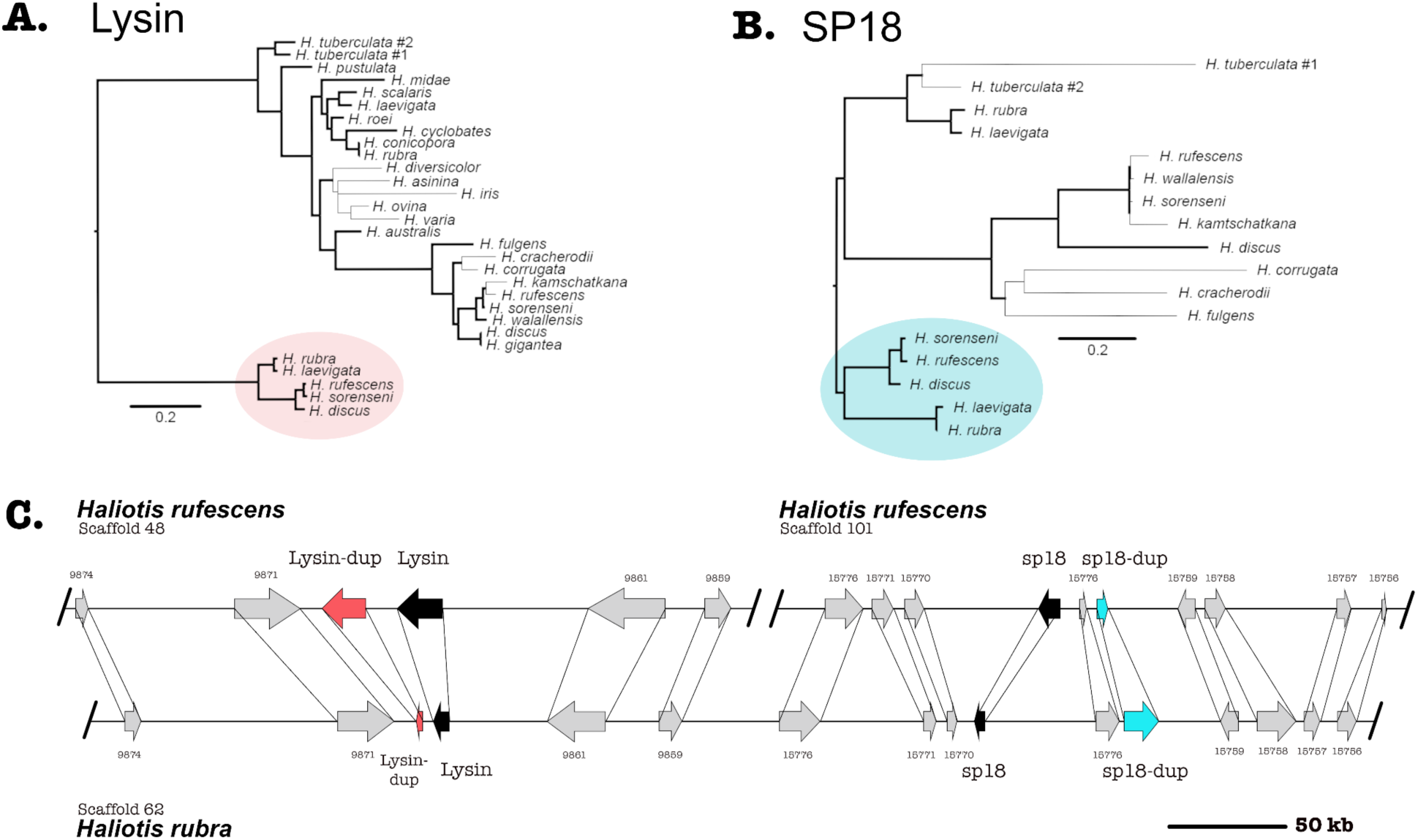
Syntenic and phylogenetic analysis indicate that four tandem acrosomal proteins are ancestral to all abalone. **A**. Ancestral duplication led to the paralogs lysin and lysin-dup (Red). Bold lines indicate greater than 80% bootstrap branch support. **B**. Ancestral duplication led to the paralogs sp18 and sp18-dup (blue). A clade specific duplication of sp18 is present in the *H. tuberculata* testes transcriptome. Bold lines indicate greater than 60% bootstrap branch support. **C**. Lysin (black), sp18 (black) and their paralogs lysin-dup (red) and sp18-dup (blue) are found in the genomes of North American and Australian abalone. Syntenic analysis indicates that paralogs are located near each other in the abalone genome, although all four paralogs are only genetically linked in Australian abalone genome assemblies. In the North American abalone H. rufescens genome assembly, lysin and sp18 are found on separate scaffolds linked with their paralogs.

Sequence analysis is consistent with the sp18-dup gene encoding a functional reproductive protein ancestral to *Haliotis*. Sp18-dup is predicted to have a signal peptide sequence and maintains a pair of cysteine residues involved in forming a structurally important disulfide-bond in sp18 (Kresge et al., 2000, 2001). Sp18-dup has not been identified in previous analysis due to the high divergence between it and sp18 (27.5% sequence identity between *H. rufescens* sp18 paralogs) obscuring homology.

Lysin-dup was identified in all abalone genomes investigated but was not detected in the testes illumina transcriptome of *H. rufescens* (Palmer et al., 2013). The absence of lysin-dup in the testes transcriptome could indicate insufficient read depth, differences in tissue-expression, or potentially pseudogenization. Since the full-length sequence of lysin-dup was not identified within the *H. rufescens* testis transcriptome, lysin sequences were used instead to identify lysin-dup exons within abalone genomes. However, divergence between lysin and lysin-dup likely prevented the identification of full-length coding sequence from abalone genomes. Only exons 2-4 could be identified (79% of query sequence) within *H. rufescens* and *H*.*rubra*. The missing exons 1 and 5 contain the signal peptide and the N- and C-termini of the molecule. In lysin, the N- and C-terminus are under strong positive selection promoting extensive divergence that reduces the ability to identify these exons using homology-based approaches (Lee et al., 1995; Lyon & Vacquier, 1999).

In the Australian abalone genomes lysin, lysin-dup, sp18, and sp18-dup are all found within a single contig with 233 kb separating the paralog pair of lysin and lysin-dup from the paralog pair of sp18 and sp18-dup. But in the genome of the North American abalone species *H. rufescens*, the paralog pair of lysin and lysin-dup are on a separate scaffold from the paralog pair of sp18 and sp18-dup. We compared the Australian contig containing the four acrosomal protein paralogs with the two *H. rufescens* contigs containing the lysin and sp18 paralog pairs respectively (**Figure 1C**). We found several ORFs surrounding each paralog pair in *H. rufescens* that were found in the same order between in *H. rubra*, indicating synteny between scaffolds. All four acrosomal proteins being located near each other in the same scaffold in the *H. rubra* genome suggests that tandem duplication led to recurrent duplications of this protein family (Reams & Roth, 2015). The sp18 ORF codes in a different direction than the other paralogs, suggesting that transposition and inversion may have also contributed to duplications within this protein family (Reams & Roth, 2015).

### Patterns of divergence of ancestral acrosomal protein paralogs indicate subfunctionalization

Lysin, sp18, and sp18-dup all contained sites detected to be subjected to positive selection. (**Table 1**). Lysin-dup did not show signatures of positive selection, though this could be due to having insufficient sequences to provide the statistical power to conduct the test (**Table 1**) (Anisimova, Bielawski, & Yang, 2001). Clustering and distribution of amino acid sites undergoing positive selection can identify regions important to the function of rapidly evolving genes (Anisimova et al., 2001). For example, many of the sites in lysin that are undergoing positive selection (11/23) are in a region of the molecule that binds its egg receptor VERL (D. B. Wilburn et al., 2018). We investigated the distribution of sites undergoing positive selection in sp18 and sp18-dup. Similar regions of the molecule undergoing positive selection in both paralogs would suggest a shared biochemical mechanism while differences in distributions of positively selected sites would indicate divergence in biochemical mechanism.

**Table 1:** Acrosomal protein paralogs are evolving under positive selection. Codon substitution models were used to analyze sequences of sp18, sp18-dup, lysin-dup, and lysin. Site models allowing for several neutral models (M1a, M7, and M8a) or selection models (M2a, M8, and M8a) allowing for variation among sites, were fit to the data using PAML. Sites undergoing positive selection were detected in sp18 and lysin for all model comparisons. A more powerful test (M8a v M8) detected positive selection in sp18-dup as well as sp18 and lysin. Estimates of the likelihood ratio statistic (**-2Δl)**, *dN/dS*, and the percentage of sites that are under positive selection are given. Significant tests are highlighted in yellow. (*, significant at P < 0.05; **, significant at P < 0.005.)

| Gene | Model | -2ΔI | $d_n/d_s$ | % Positively Selected Sites |
| --- | --- | --- | --- | --- |
| Sp18 | M1a vs M2a | 84.2** | 4.4 | 48 |
|  | M7 vs M8 | 95.3** | 4.2 | 48 |
|  | M8a v M8 | 84.3** | 4.2 | 48 |
| Sp18-sup (do you man “dup”?) | M1a vs M2a | 4.1 | - | - |
|  | M7 vs M8 | 4.1 | - | - |
|  | M8a v M8 | 4.1* | 1.8 | 49 |
| Lysin-dup | M1a vs M2a | 1.3 | - | - |
|  | M7 vs M8 | 1.5 | - | - |
|  | M8a v M8 | 1.3 | - | - |
| Lysin | M1a vs M2a | 156.1** | 1.2 | 21 |
|  | M7 vs M8 | 156.7** | 1.1 | 22 |
|  | M8a v M8 | 141.1** | 1.1 | 22 |

By mapping sites under positive selection onto a protein alignment of sp18 and sp18-dup, we determined that sites under positive selection in either paralog are non-randomly distributed across the protein alignment and differentially clustered. We analyzed the selected sites in the primary sequence alignment with a parametric adaptation of the runs test (Wald-Wolfowitz test) (**Figure 2A**) (Magel, 1997). This analysis showed that there were significant runs of sites undergoing positive selection in either paralog (p-value = 0.026), consistent with different regions evolving under positive selection among paralogs. Positive selection acting on different regions of the protein alignment is consistent with subfunctionalization of paralogs.

**Figure 2:**
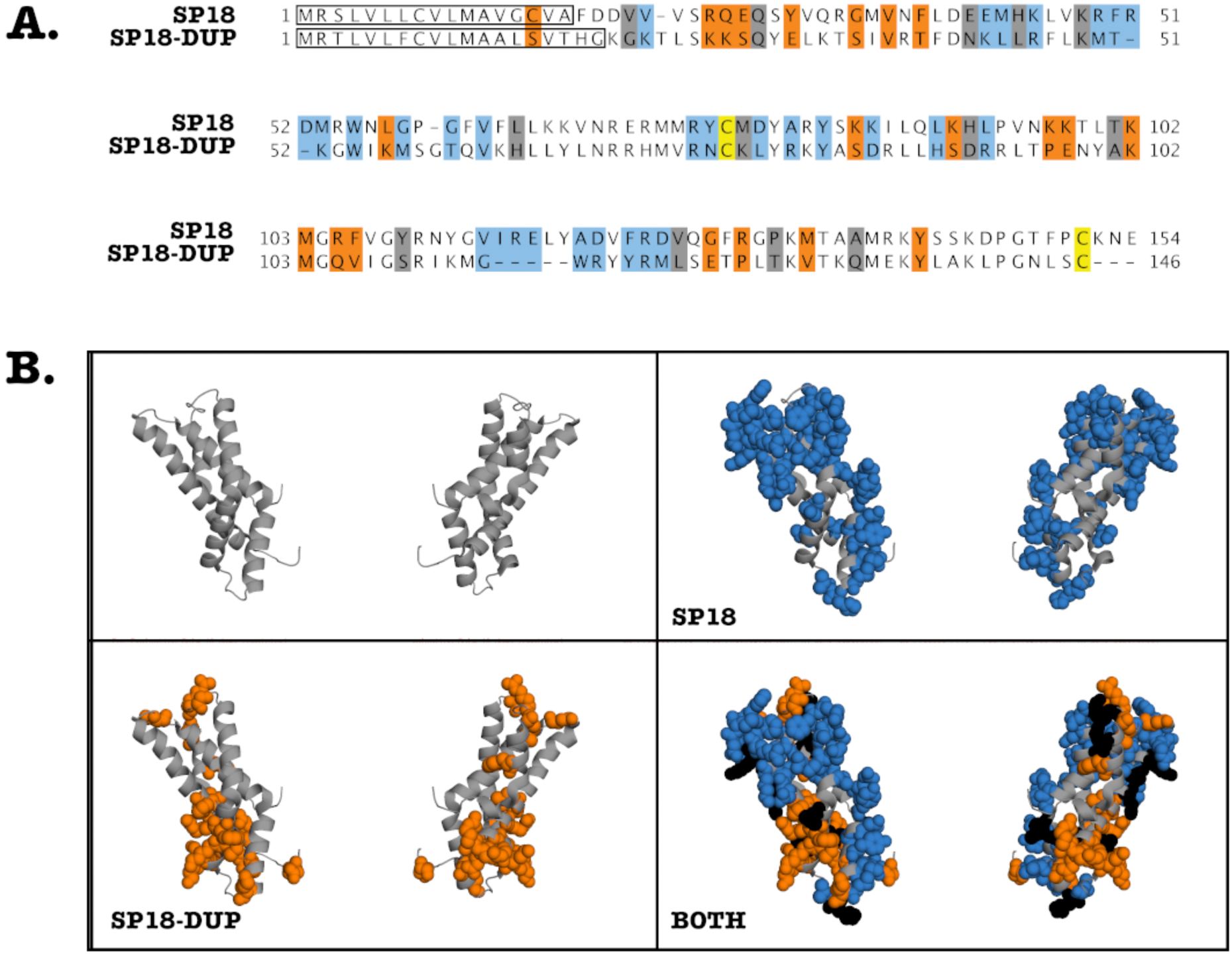
Clustering of positively selected sites is consistent with subfunctionalization of SP18 and SP18-dup. **A**. Alignment of *H. fulgens* sp18 and *H. sorenseni* sp18-dup mature protein sequences. Sites under positive selection are highlighted. Blue (Sp18), Orange (Sp18-dup), and Black (both). A modified parametric runs test was used to determine that there were statistically significant runs in linear sequence space of sites under positive selection in sp18 or sp18-dup (p-value - 0.0XX). **B**. Sites under positive selection in sp18 or sp18-dup were mapped onto the crystal structure of sp18 from *H. fulgens* and appear clustered in 3-D space.

We investigated clustering of positively selected sites in three-dimensional space. By mapping sp18 and sp18-dup positively selected sites onto the sp18 structure, it is visually apparent that there are distinct clusters of sites under selection between paralogs (**Figure 2B**). Using the plane of best fit through the crystal structure we divided the molecule into “left” and “right” sides agnostic to the location of positively selected sites. To define the “top” and “bottom” of the molecule we used a plane perpendicular to the plane of best fit. Sites under positive selection in sp18-dup were statistically more likely to be on the “right” side than on the “left” (p-value <0.05) (**Supplementary Figure 2**). Sp18 sites, using the same test, showed no statistically significant difference from the null distribution. However, sites under positive selection in sp18 were enriched on the “top” of the molecule rather than the “bottom” (p-value <0.05) while the distribution of sp18-dup positively selected sites did not significantly differ (Supplementary Figure 2). These two tests show that sites under selection in sp18 and sp18-dup are distributed differently across their three-dimensional structures.

We also developed a test to examine whether sites under selection in sp18 and sp18-dup were statistically more likely to be adjacent to a site under selection from the same paralog. Such a pattern of clustering would indicate a spatial relationship between positively selected sites belonging to a particular paralog. For each site under positive selection in sp18 or sp18-dup, we identified whether the closest positively selected site in three-dimensional space was significantly more likely to belong to the same paralog. We found that sites under selection in both paralogs were more likely to have the closest positively selected site belong to the same paralog rather than the other paralog according to a chi-squared test (p-value < 0.01). Together, the runs test analysis and three-dimensional analyses point to diversifying selection post-duplication of these proteins to promote functional diversification.

Because Lysin-dup was not detected to be under positive selection, we did not test for differences in sites under selection between lysin paralogs. However, we did evaluate how the lysin-dup sequence diverged from lysin. Many of the sites shown to be undergoing positive selection in lysin differ in sequence from lysin-dup (13/14) when comparing the *H. rufescens* sequences (**Figure 3B**). Although this comparison is not significant (p = 0.088), this suggests similar sites driving the diversification in sequence of lysin between species and between lysin and its paralog lysin-dup.

**Figure 3.**
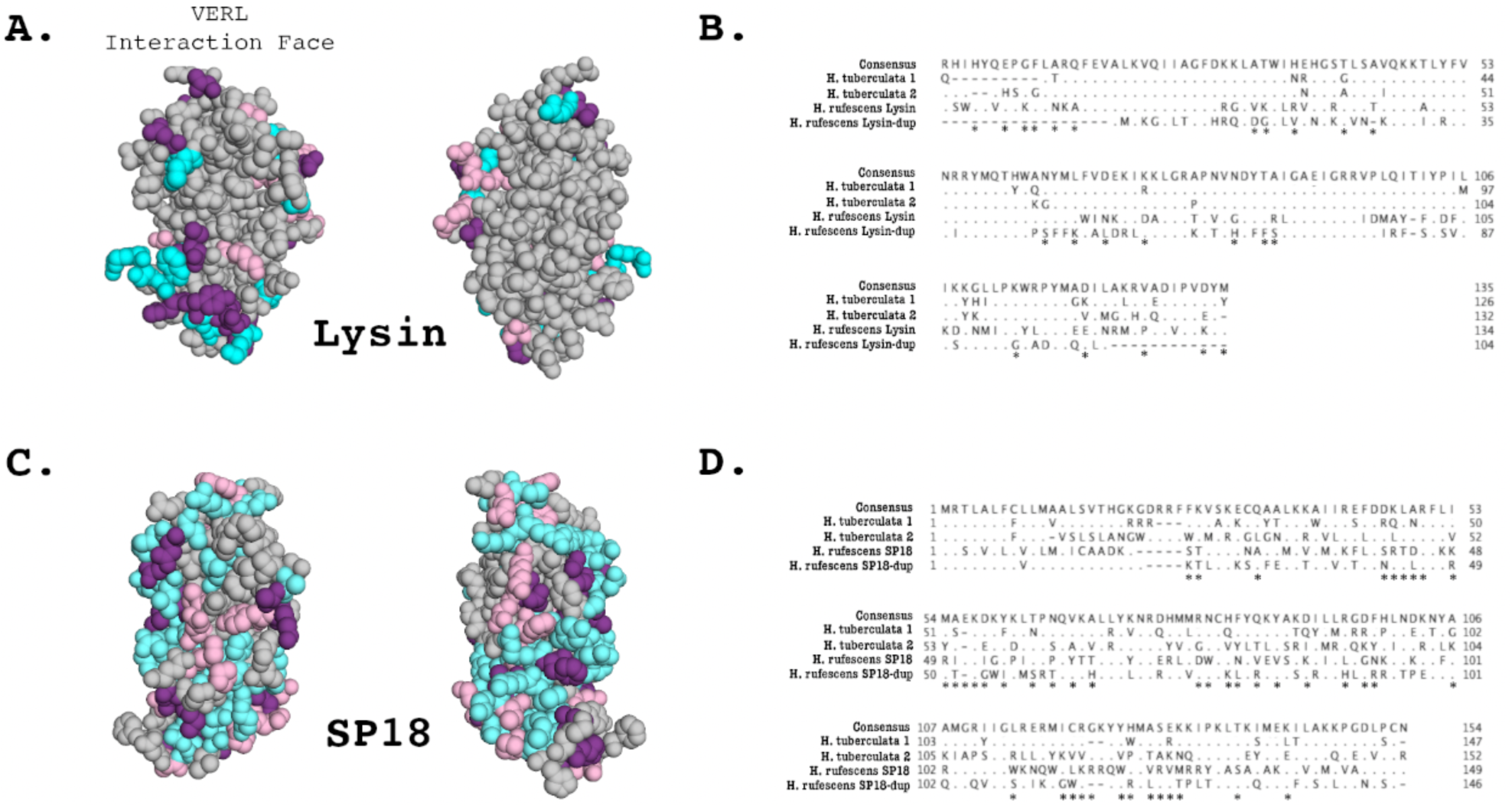
Duplication and divergence of lysin and sp18 paralogs in *H. tuberculata*. **A**.) Sites that differentiate *H. tuberculata* lysin paralogs from *H. rufescens* lysin are mapped onto the *H. rufescens* NMR structure (PDB 5UTG). Blue – Sites that differ between *H. rufescens* lysin and *H. tuberculata* Copy #1, Pink – Sites that differ between *H. rufescens* lysin and *H. tuberculata* Copy #2, Purple – Sites that differ between *H. rufescens* lysin and both *H. tuberculata* lysin paralogs. Notably the face of the lysin molecule that interacts with VERL harbors most substitutions between *H. tuberculata* paralogs, suggestive of a shared interaction face for both *H. tuberculata* lysin paralogs. **B**.) Alignment of Lysin paralogs in *H. rufescens* and *H. tuberculata*. Both sets of paralogs arose from independent duplication events. Asterisks indicate sites that are shown to be under positive selection in *H. rufescens* lysin. **C**.) Sites that differ between *H. tuberculata* sp18 paralogs are mapped onto the crystal structure of sp18 *from H. fulgens* (PDB 1GAK). Blue – Sites that differ between *H. rufescens* sp18 and *H. tuberculata* Copy #1, Pink – Sites that differ between *H. rufescens* sp18 and H. tuberculata Copy #2, Purple – Sites that differ between *H. rufescens* sp18 and both H. tuberculata sp18 paralogs. **D**.) Alignment of sp18 paralogs in *H. rufescens* and *H. tuberculata*. Both sets of paralogs arose from independent duplication events. Asterisks indicate sites that are shown to be under positive selection in *H. rufescens* sp18.

### *H. tuberculata* sp18 and lysin duplications are species-specific

Previous work described a lysin duplication unique to *H. tuberculata* (Clark et al., 2007). The lysin paralogs were shown to be evolving under positive selection and to be maintained in the testis proteome (Clark et al., 2007). Sites that vary between European lysin paralogs are largely located on the face of the molecules interacting with lysin receptor VERL (**Figure 3A**). To investigate the presence of additional sp18 and lysin paralogs in *H. tuberculata*, we constructed a long-read PacBio testis transcriptome. Performing tBLASTN searches of the *H. tuberculata* transcriptome for lysin and sp18 revealed the previously described species-specific duplication of lysin (*H. tuberculata* lysin copy #1 and copy #2) and a novel duplication of sp18 (*H. tuberculata* sp18 copy #1 and copy #2).

Phylogenetic analysis indicates the *H. tuberculata* sp18 paralogs are the result of a recent duplication and not ancestral to *Haliotis*. Sequence information from other abalone species is needed to determine whether this duplication is species-specific to *H. tuberculata*; it appears to be specific to the abalone clade containing the European species. The signal sequences of the *H. tuberculata* sp18 paralogs are more similar to each other than to signal sequences from other species’ sp18 paralogs. Signal sequences are not part of the mature protein and not subjected to the same evolutionary pressures driving rapid divergence, therefore these sequences show more conservation between closely related paralogs. This similarity in signal sequence between *H. tuberculata* sp18 paralogs (15/19 sites are identical) further supports that these paralogs are the result of a non-ancestral duplication. Despite being a more recent duplication of sp18, the paralogs have a low sequence identity (39%), lower than that of the *H. tuberculata* lysin paralogs (83%). This is consistent with sp18 having a higher *d*_*N*_*/d*_*S*_ and evolving more rapidly than lysin (**Table 1**). *H. tuberculata* sp18 paralogs maintain a pair of structurally important cysteine residues involved in forming a disulfide bond. Rapid sequence divergence, no premature stop codons, and both genes being expressed in the testis transcriptome are all indicators that both sp18 paralogs (referred to here as *H. tuberculata* copy #1 and copy #2) are likely to be functional.

The *H. tuberculata* sp18 copy #1 is the more divergent to the ancestral sequence than copy #2, as indicated by its long branch in the sp18 phylogeny (**Figure 1B**). When comparing the sequence identity of *H. tuberculata* sp18 paralogs to *H. rubra* sp18 (an outgroup sequence), copy #1 shows a lower sequence identity (41%) than copy #2 (69%). This rapid sequence divergence of copy #1 without accruing mutations causing pseudogenization suggests strong positive selection. However pairwise *d*_*N*_*/d*_*S*_ between *H. tuberculata* sp18 paralogs could not be reliably estimated due to extensive divergence resulting in saturation (multiple substitutions per site) (Swanson & Vacquier, 1995). Maintenance of both paralogs in the testis proteome despite the observed sequence divergence would indicate that both paralogs are being selected for functions, presumably related to fertilization. We used data dependent acquisition mass spectrometry to identify peptides belonging to either paralog in the *H. tuberculata* testes proteome. Diagnostic peptides were detected for copy #1 but not copy #2. Despite being the more divergent sp18 sequence, copy #1 is maintained in the proteome. This result indicates that copy #1 is likely important for fulfilling sp18’s membrane fusion function in *H. tuberculata*. \

### Lack of Recent Duplications of Egg Coat Proteins

We generated an ovary PacBio transcriptome for *H. tuberculata* to identify VEZP proteins. Using the 33 VEZP and ZP-domain sequences from the *H. rufescens* ovary transcriptome as the initial query sequences, exhaustive tBlastn searches of the ovary transcriptome were used to identify all cDNA sequences with sequence similarity to any *H. rufescens* VEZP. ZP module protein sequences were extracted from our *H. tuberculata* cDNA hits and the 33 *H. rufescens* VEZPs and then were aligned to construct a phylogeny (**Figure 4**). Clustering of *H. tuberculata* and *H. rufescens* ZP module sequences indicate that these distantly related abalone species have the same complement of ZP-proteins in their transcriptomes. In *H. tuberc*ulata’s ovary transcriptome, orthologs of 32 of the 33 *H. rufescens* ovary ZP-domain proteins were identified. No VEZPs, including VERL and its most closely related paralogs VEZP14 and VEZP9 were duplicated. The only missing sequence belonged to ZPC, a gene whose cDNA sequence contains a premature stop codon in *H. rufescens* and for which no peptides were detected in the *H. rufescens* VE proteome (Aagaard et al., 2010). Therefore, ZPC is likely pseudogenized in *H. rufescens* and its expression its expression is no longer maintained in European abalone. Remarkably, no new ZP-module-containing proteins were identified in *H. tuberculata* despite the species having multiple clade-specific duplications of acrosomal proteins. These results suggest that the clade-specific maintenance of duplicated sperm acrosomal proteins found in the European abalone *H. tuberculata* are unlikely to be the result of duplicated egg proteins.

**Figure 4.**
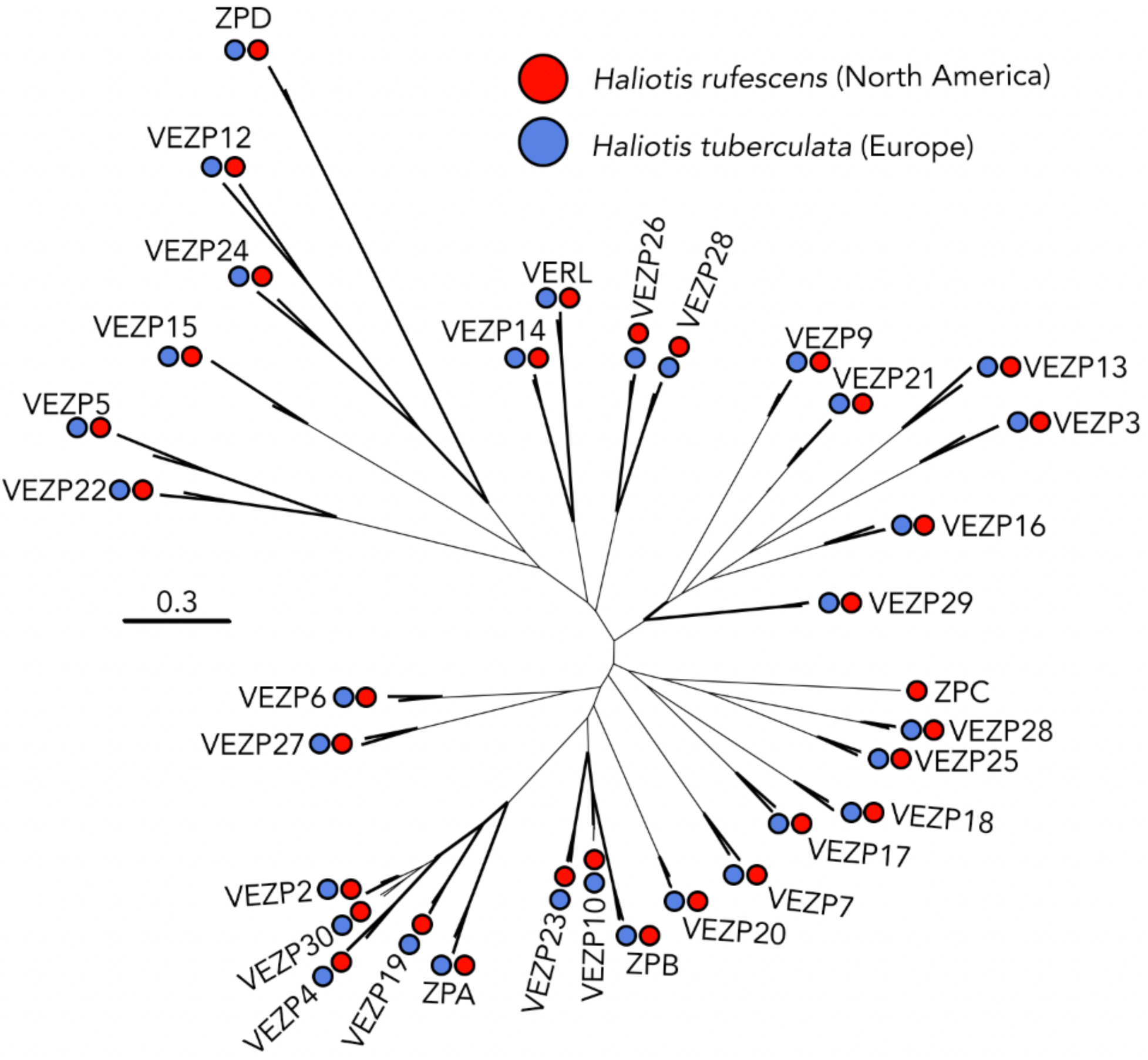
VEZP proteins are conserved across abalone species. The distantly related abalone species *H. rufescens* (Red) and *H. tuberculata* (Blue) share the same vitelline envelope zona pellucida (VEZP) proteins within their ovary transcriptomes. Although there are species-specific duplications of lysin and sp18 *in H. tuberculata*, there is no evidence of species-specific duplications of VERL or other VEZP proteins. Bold lines indicate greater than 80% bootstrap branch support.

## Discussion

Despite decades of research examining the evolution of abalone fertilization genes, only recently have genomic resources been available that enable a broad investigation into the evolution of the protein families to which lysin and VERL belong. Here, we explore the contributions of duplication and sequence divergence to the evolution of abalone fertilization genes across the genus *Haliotis*. For our investigation we generated ovary and testes transcriptomes from the European abalone *H. tuberculata* and utilized recently published North American and Australian abalone genomes and a North American abalone testes transcriptome (Palmer, McDowall et al. 2013, Nam, Kwak et al. 2017, Botwright, Zhao et al. 2019, Gan, Tan et al. 2019, Masonbrink, Purcell et al. 2019). We discovered novel duplications of both lysin and sp18 ancestral to abalone, indicating that abalone lysin and sp18 are members of an ancestral abalone protein family with four members. The newly discovered sp18 paralog (sp18-dup) was shown to be undergoing positive selection, like lysin and sp18, and expressed in the testes of North American abalone. Further, differences in clustering of positively selected sites in sp18 compared to sp18-dup is consistent with models of subfunctionalization. We investigated whether there are clade-specific duplications of abalone VEZPs or acrosomal proteins. In addition to a species-specific lysin duplication described in a previous publication (Clark et al., 2007), the *H. tuberculata* testes transcriptome contains a clade-specific duplication of sp18 not found in Australian or North American abalone species. However, no duplications of VERL or other VEZPs were observed between North American or European abalone, indicating that VEZP gene content is conserved across the genus *Haliotis*. Together, this data demonstrates that recurrent duplication and diversification driven by positive selection drives the evolution of an acrosomal protein family involved in fertilization in *Haliotis*.

### Recurrent Duplication and Positive Selection of Acrosomal Proteins in Abalone

In the *H. rubra* abalone genome, the paralogs lysin, lysin-dup, sp18, and sp18-dup are found on a single scaffold. This clustering within the genome indicates that ancestral tandem duplication events occurred leading to the creation of this acrosomal protein family (Reams & Roth, 2015). Further, three of the four ancestral paralogs were shown to be maintained in the testis transcriptome and to be evolving under positive selection, a common characteristic of reproductive proteins.

This evolutionary pattern of duplication paired with sequence diversification found in the abalone acrosomal protein family can be compared to protein families in other taxa which contain sperm proteins mediating fertilization. Notably, the mammalian Izumo gene family contains four ancestral paralogs whose members all show testes-specific tissue expression in humans (Grayson & Civetta, 2012). Izumo1 is an essential gene for sperm-egg plasma membrane fusion in mammals that functions by binding the egg plasma membrane protein JUNO (Bianchi et al., 2014). There is evidence that the other three Izumo paralogs may also possess important, although potentially varied, functions in fertility (Ellerman et al., 2009). All four paralogs have been shown to be undergoing rapid sequence evolution in at least one mammalian lineage, for Izumo1, 2, and 3 this is driven by positive selection and for Izumo4 this appears to be driven by relaxed selection (Grayson, 2015; Grayson & Civetta, 2012). Given that both the abalone acrosomal protein family and the mammalian Izumo family both contain multiple paralogs showing testis-specific function, subfunctionalization may be a common driver of the evolution of fertilization and reproductive genes across taxa. Understanding how fertilization proteins emerge and evolve can be important for identifying and understanding mechanisms of fertilization across diverse taxa.

### Recurrent Subfunctionalization of Abalone Acrosomal Proteins

Differences in optimal mating rates for sperm and eggs can drive antagonistic coevolution of reproductive proteins. Under this sexual conflict scenario, evolution of egg coat proteins interacting with sperm acrosomal proteins could lead to constrained evolution on the sperm side (Gavrilets & Waxman, 2002). Duplication followed by diversification of sperm fertilization proteins can be an important means of sperm escaping evolutionary constraints imposed by egg protein evolution. For two duplication events within the abalone acrosomal protein family there is evidence of subfunctionalization from either functional experiments (lysin vs. sp18, (Kresge et al., 2001; Swanson & Vacquier, 1997) or site-clustering analysis (sp18 vs. sp18-dup, current manuscript).

Plasma membrane fusion in fertilization or other contexts is traditionally thought to consist of two steps, binding and fusion (Bianchi & Wright, 2020). In sea urchins, both steps are mediated by different regions of the same protein, su-bindin (Ulrich, Otter, Glabe, & Hoekstra, 1998; Vacquier & Moy, 1977; Vacquier & Swanson, 2011). However, in abalone these steps may have been partitioned between sp18 and sp18-dup via subfunctionalization. Abalone eggs have a thin layer directly overlaying the surface of the plasma membrane which morphologically resembles a duplication of the elevated VE (Mozingo et al., 1995). Just as lysin binds the VE protein VERL, sp18 may bind a VEZP protein found within the thin layer overlaying the abalone egg plasma membrane. Indeed, in addition, to having a strong fusagenic function, sp18 has been demonstrated to bind to VEZP proteins, an unsurprising trait for a lysin paralog (Aagaard et al., 2010). One possibility is that the subfunctionalization of sp18 and sp18-dup may have been driven by the separation of the steps of plasma membrane binding and fusion between paralogs.

While the paralogs lysin-dup and lysin do show high sequence divergence, only lysin shows evidence of positive selection. Therefore, the observed sequence divergence is likely driven by lysin’s evolution post-duplication. Unlike the other acrosomal protein family paralogs discussed in this paper, lysin-dup is not detected in the testis transcriptome. However, there is an appealing hypothesis as to its potential function. In abalone egg coats there are two VEZP proteins capable of binding lysin, VERL the major binding partner of lysin and VEZP-14 the most recent paralog of VERL (Aagaard et al., 2013). It is possible that lysin-dup may be the binding partner of VEZP-14 and if true this could explain why lysin shows correlated evolution with VERL but not VEZP-14 (Aagaard et al., 2013). Currently there is insufficient data to test for correlated rates of evolution between lysin-dup and VEZP-14. However, further molecular and biochemical characterization through binding kinetic analysis could test the hypothesis that lysin-dup and VEZP-14 interact.

### Species-Specific Duplications of Acrosomal Proteins in Abalone

Previous work described a lysin duplication unique to *H. tuberculata* and maintained in the testis proteome (Clark et al., 2007). In this study a clade-specific duplication of sp18 was discovered within the *H. tuberculata* transcriptome. Despite having two acrosomal protein duplications, European abalone’s ovary transcriptome did not reveal any novel VEZP protein sequences indicative of a duplication event. While gene duplications are an ongoing contributor to the evolution of sperm fertilization genes in abalone, this may not be true for egg fertilization genes. Our data suggests that it is not duplications on the egg side driving the duplication of abalone acrosomal proteins in *H. tuberculata*. This could be explained by different selective pressures on the sperm and the egg, such as sperm competition and polyspermy risk (Carlisle & Swanson, 2020). Further, It does not seem that a process of gene birth and loss explains the evolution of abalone’s acrosomal protein family since all paralogs are maintained in the transcriptome and have accrued no pseudogenizing mutations. A hypothesis for the duplication and diversification of acrosomal protein paralogs in *H. tuberculata* is that paralogs are specialized for different binding sites of their egg receptor or different allelic variants of their receptor. For example, *H. rufescens* VERL has 22 tandem ZP-N domains with three unique amino acid sequences, *H. tuberculata* VERL may show similar differences in ZP-N sequences and *H. tuberculata* lysin paralogs may be optimized for binding different ZP-N sequences (Galindo, Moy, Swanson, & Vacquier, 2002). In addition, the abalone *H. tuberculata* VERL may be polymorphic, as seen for *H. corrugata* VERL, and lysin paralogs are optimized for VERL allelic variants (Clark et al., 2009). This study observed that sites that vary between *H. tuberculata* lysin paralogs are largely located on the face of the molecules interacting with lysin receptor VERL (**Figure 3A**). Unlike the distribution of positively selected sites between sp18 and sp18-dup where sites are differentially clustered on the protein structure. This pattern of diversification may be suggestive of specialization of function, such as interacting with different VERL allelic variants or VERL ZP-N domains. Further characterization of VERL in *H. tuberculata* and population-level variation is necessary to explore these hypotheses.

## Conclusion

This study characterizes duplication events of a sperm acrosomal protein family with functions directly associated with fertilization. Although lysin was one of the first fertilization proteins discovered and the first for which an egg binding partner was defined, its evolutionary origins are unknown. By placing duplication events of lysin and sp18 within their genomic context and identifying clade-specific duplication events, this study has revealed the importance of duplication for the evolution of this protein family that has previously been unknown. We describe six acrosomal protein paralogs arising from both ancestral and clade-specific duplication events (**Figure 5**). Recurrent duplication events of sperm acrosomal proteins have occurred throughout the evolutionary history of abalone. For the two abalone species with transcriptomic data both have paralogs maintained in the testis transcriptome. Remarkably none of these genes have been pseudogenized and many are undergoing strong positive selection consistent with maintenance of their function in abalone reproduction. Further inquiry is required to investigate why these proteins are undergoing duplication, the functional consequences of these duplication events, and whether other fertilization proteins in other species (as also seen for the mammalian Izumo family) are undergoing recurrent duplication events.

**Figure 5:**
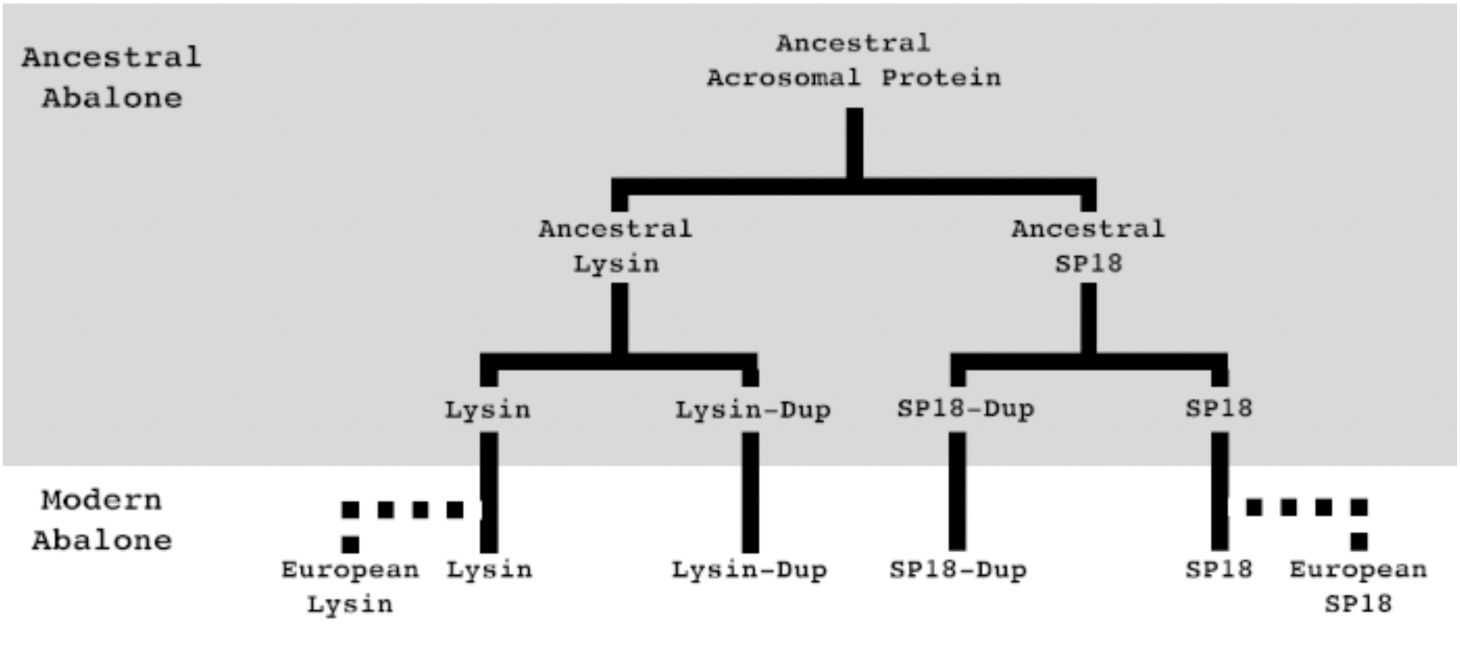
Summary of abalone acrosomal duplications. Abalone acrosomal proteins have duplicated multiple times over the course of abalone evolution. Five duplication events can describe the currently identified protein family. Most duplications are ancestral to all abalone, but species-specific duplications in European abalone point to duplication paired with positive selection still playing an important role in the evolution of abalone fertilization pathways. Solid lines indicate orthologs present in all abalone, dotted lines indicate species-specific paralogs.

## Author Contributions

JAC and WJS designed the research. JAC and MG performed the research. JAC and WJS analyzed the data. JAC wrote the paper.

## Acknowledgements

This study was supported by NIH Grant HD076862 to Willie J. Swanson and a National Science Foundation Graduate Research Fellowship to Jolie A. Carlisle. Megan Glenski was supported by University of Washington School of Medicine-Gonzaga University Regional Health Partnership. Thank you to France Haliotis for help acquiring *H. tuberculata* samples, to Evan Cox, Dr. Daniel Promislow, Dr. Josh Schraiber, and Dr. Damien Wilburn for help with data analysis, and to Alberto Rivera, Dr. Jan Aagaard, and Dr. Bryce Taylor for useful discussions and comments. All research was performed on the traditional lands of the Duwamish Tribe. To learn more about the Duwamish Tribe and their continuing legacy, please visit https://www.duwamishtribe.org/.

---

**Supplementary Figure 1:**
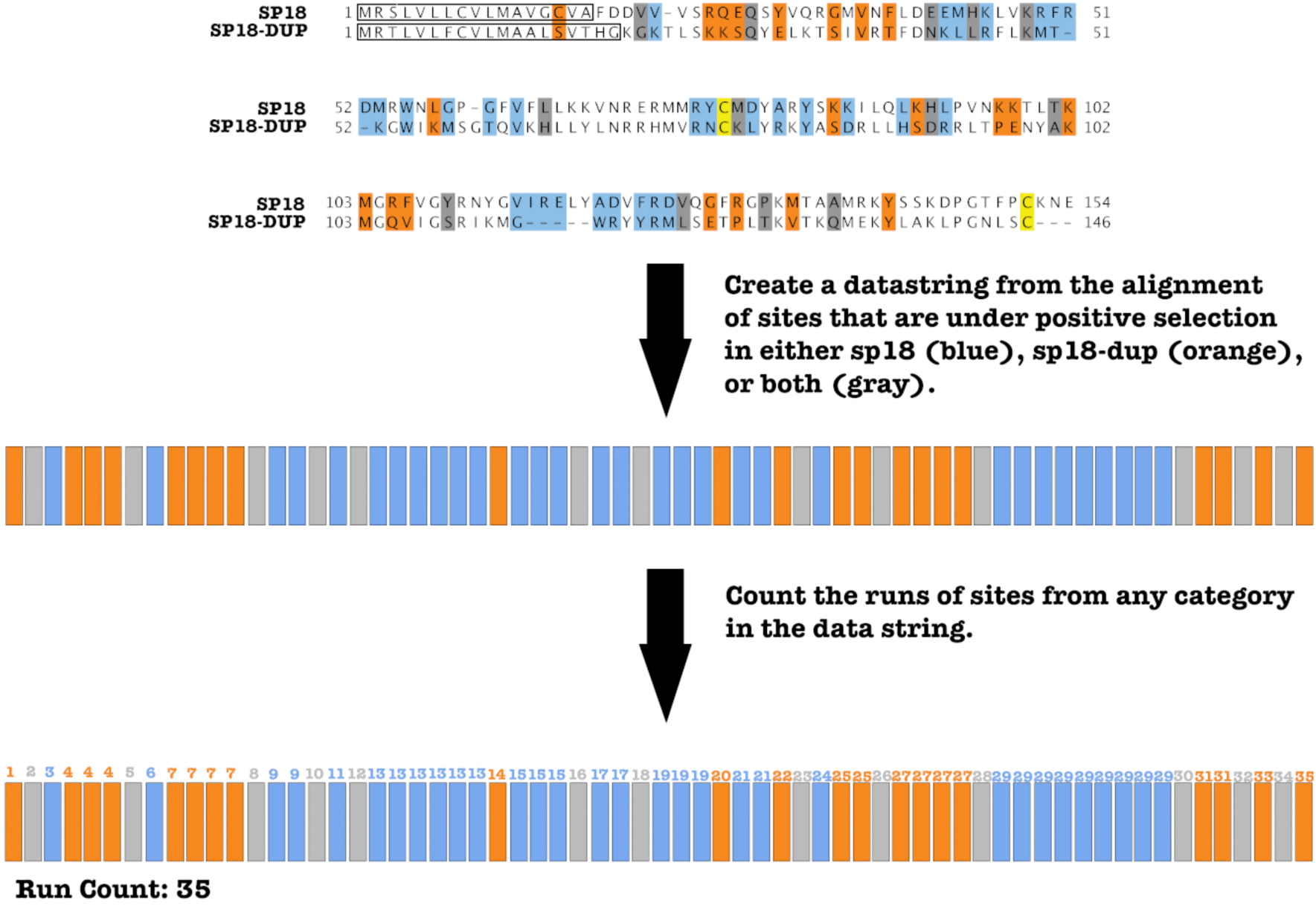
Calculation of Runs of Positively Selected Sites. The Wald-Wolfowitz runs test determines the randomness of a two-category data string by examining changes between categories by counting “runs.” We designed a parametric version of the test to allow the inclusion of three categories. The categories were sites under positive selection in sp18, sites under selection in sp18-dup, and sites under selection in both. The order in which these sites under selection in the categories appeared in an alignment of *H. fulgens* sp18 and *H. sorenseni* sp18-dup became our data string. For the data string generated from our paralog alignment we counted how many times the identity of sites in the string changed plus one. A visualization of the pipeline for preparing this data string and counting “runs” <u>is</u> shown here. We also performed an alternate version of this analysis where the gray sites (both) were removed when calculation changes in categories.

**Supplementary Figure 2:**
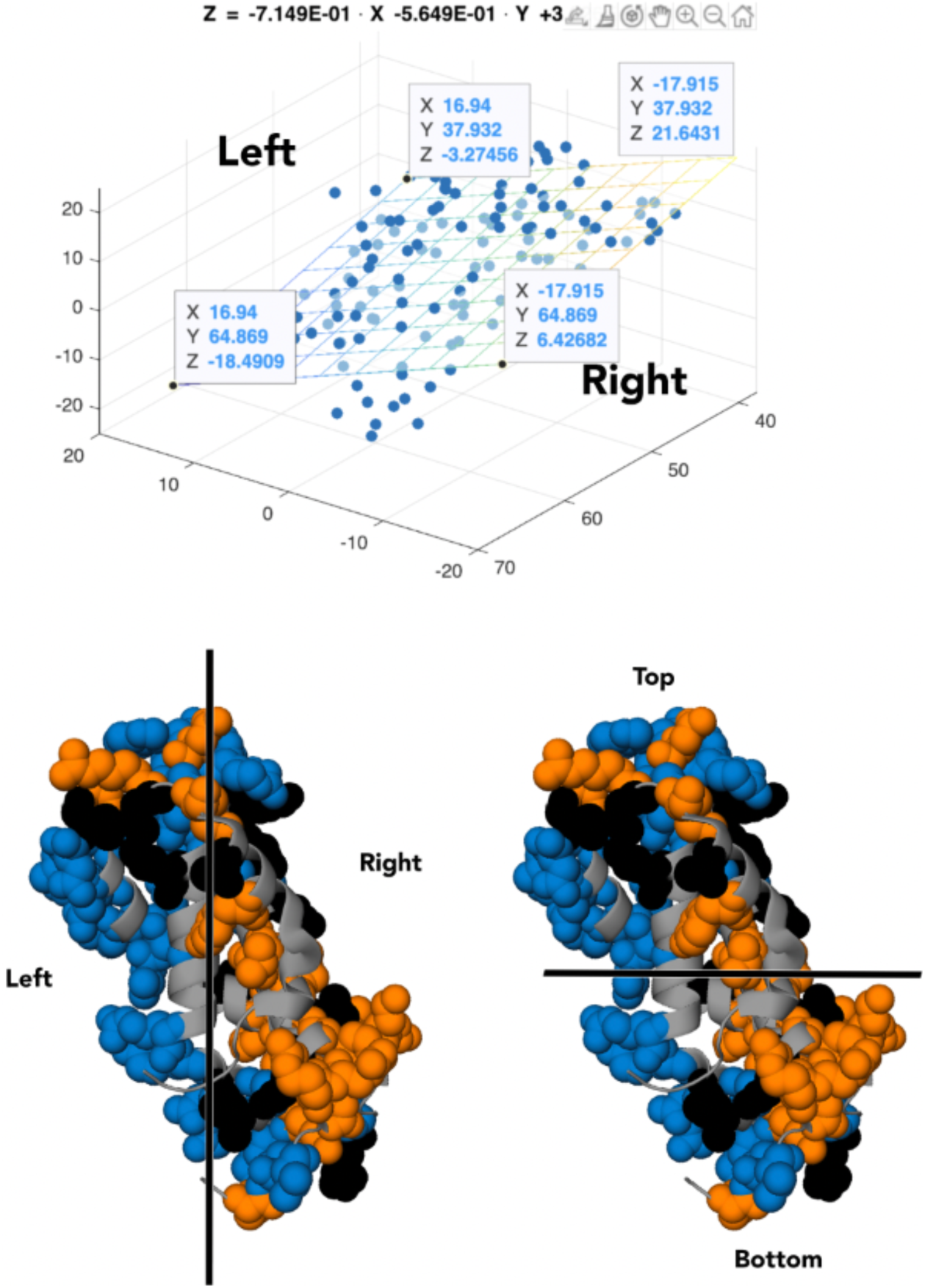
Planes separating sp18 crystal structure. **A**.) Matlab was used to calculate the plane of best fit (Supplementary File 1) through the carbon alphas of the crystal structure (PDB 1GAK). **B**.) The plane of best fit divides the sp18 crystal structure into “left” and “right” sides. **C**.) A plane perpendicular to the plane of best fit divides the sp18 crystal structure into “top” and “bottom” sides.

